# The expression of *PDCD1* and *CD274* in T cells and macrophages correlated positively with COVID-19 severity

**DOI:** 10.1101/2020.11.17.378992

**Authors:** Qianqian Gao, Shang Liu, Renpeng Ding, Huanyi Chen, Xuan Dong, Jiarui Xie, Yijian Li, Lei Chen, Huan Liu, Feng Mu

**Affiliations:** BGI-Shenzhen, Shenzhen 518083, China; Guangdong Provincial Key Laboratory of Human Disease Genomics, Shenzhen Key Laboratory of Genomics; BGI Education Center, University of Chinese Academy of Sciences, Shenzhen 518083, China; MGI, BGI-Shenzhen, China

**Author notes:** Corresponding author: Qianqian Gao.

## Abstract

The immune responses underlying the infection of severe acute respiratory syndrome coronavirus 2 (SARS-CoV-2) remain unclear. To help understand the pathology of coronavirus disease 2019 (COVID-19) pandemics, public data were analyzed and the expression of *PDCD1* (encoding PD-1) and *CD274* (encoding PD-L1) in T cells and macrophages were identified to correlate positively with COVID-19 severity.

## Introduction

COVID-19 has led to the global pandemic and infected millions of people worldwide [1]. It’s urgent to better understand the pathophysiology of the infection caused by SARS-CoV-2 [2–4]. The PD-1/PD-L1 signaling plays an essential role not only in regulating tumor immune responses but also in balancing homeostasis and tolerance in virus infection [5], but it’s role in COVID-19 is currently unclear. Thus, it is necessary to investigate how PD-1/PD-L1 signaling works during COVID-19 progress in order to deal with it.

Single-cell immune profiling was explored in COVID-19 patients using the public data, which is essential for understanding the potential mechanisms underlying COVID-19 pathogenesis.

## Methods

### Data Sets and Processing

We downloaded the gene expression matrix from GEO database (GSE145926 and GSE128033). This dataset contained 63734 cells from four healthy people, three patients with moderate infection by bilateral pneumonia and seven patients with severe infection. The detailed clinical information of patients can be obtained in the reference [6]. Single cell clustering and cell annotation was performed as previously described [6].

### Statistical analysis

The Student’s t-test (t.test in R 3.5.1) was used across three experiment designs. *P<0.05, **P<0.005, ***P< 0.001. Values are presented as mean Standard deviation (SD). Error bars represented the SD.

## Results

### Single-cell analysis revealed distinct immune cell subsets

Publicly available data of bronchoalveolar cells from three moderate (M1-M3) and six severe (S1-S6) COVID-19 patients, and four healthy controls (HC1-HC4) were collected for analysis (66630 cells, Table S1) [6]. 31 clusters were identified by classical signature genes according to the reference (Fig. 1) [6]. CD68-Macrophage (0, 1, 2, 3, 5, 6, 7, 8, 9, 11, 12, 15, 19, 21, 24, 30) dominated among these populations, followed by CD3D-T cell (4, 10, 16).

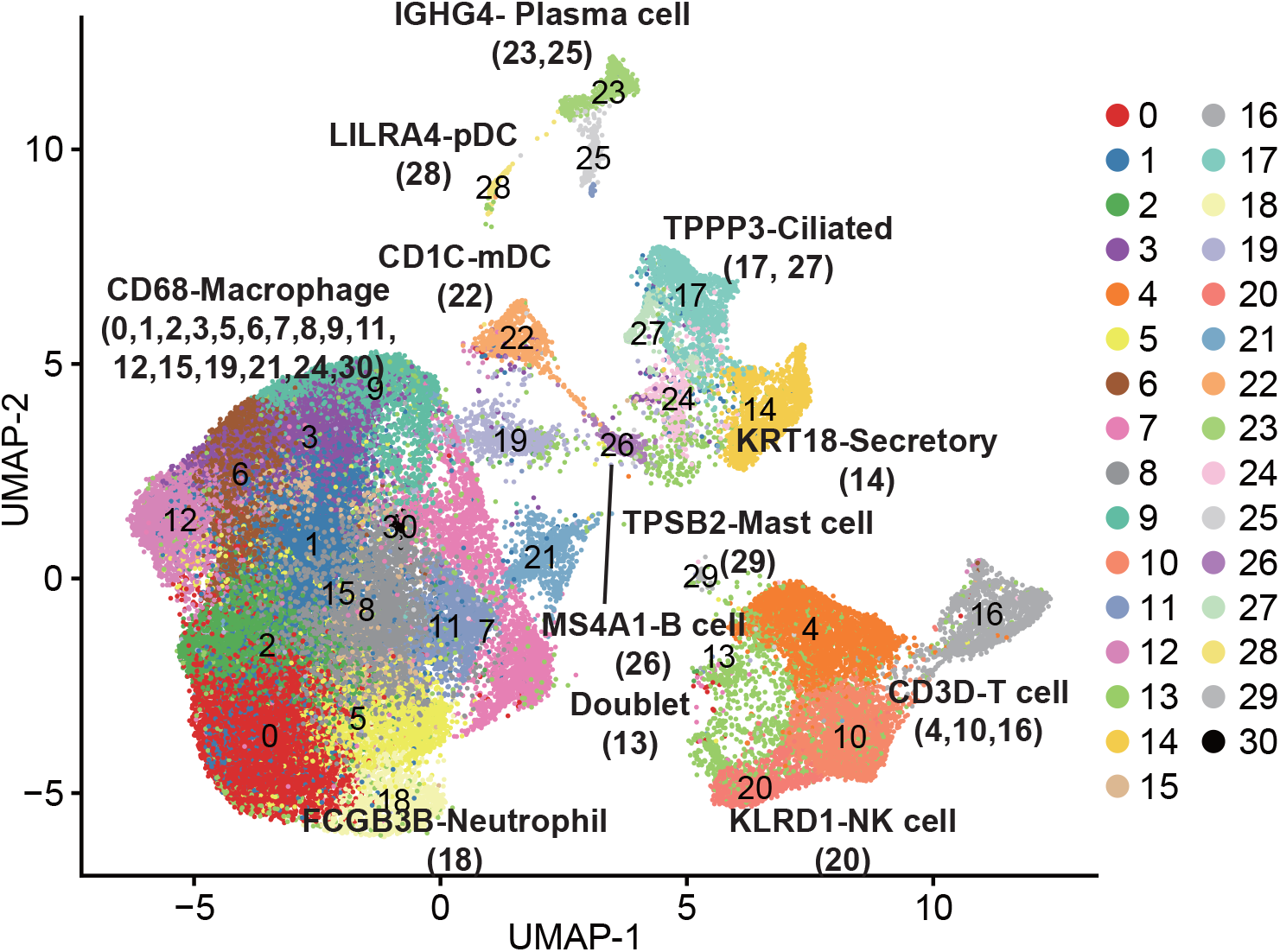

### Various expression patterns of *PDCD1* and *CD274* in different cell subpopulations

Expression of *PDCD1* was first analyzed at the patient group level in different cell subpopulations, and four trends of their expression dynamics were observed (Fig. 2). *PDCD1* expression was gradually elevated in T cell, B cell, myeloid dendritic cells (mDCs), and macrophages from HC to mild cases then to severe patients (Fig. 2A). In the 2^nd^ trend, *PDCD1* expression was specifically increased in plasma cells and epithelial cells in severe patients but not in mild patients (Fig. 2B). For the 3^rd^ trend, *PDCD1* expression was upregulated in mild patients but slightly reduced in severe patients in NK and plasmacytoid dendritic cells (pDCs) (Fig. 2C). No expression of *PDCD1* was detected in mast cells and neutrophils in the 4^th^ trend (Fig. 2D).

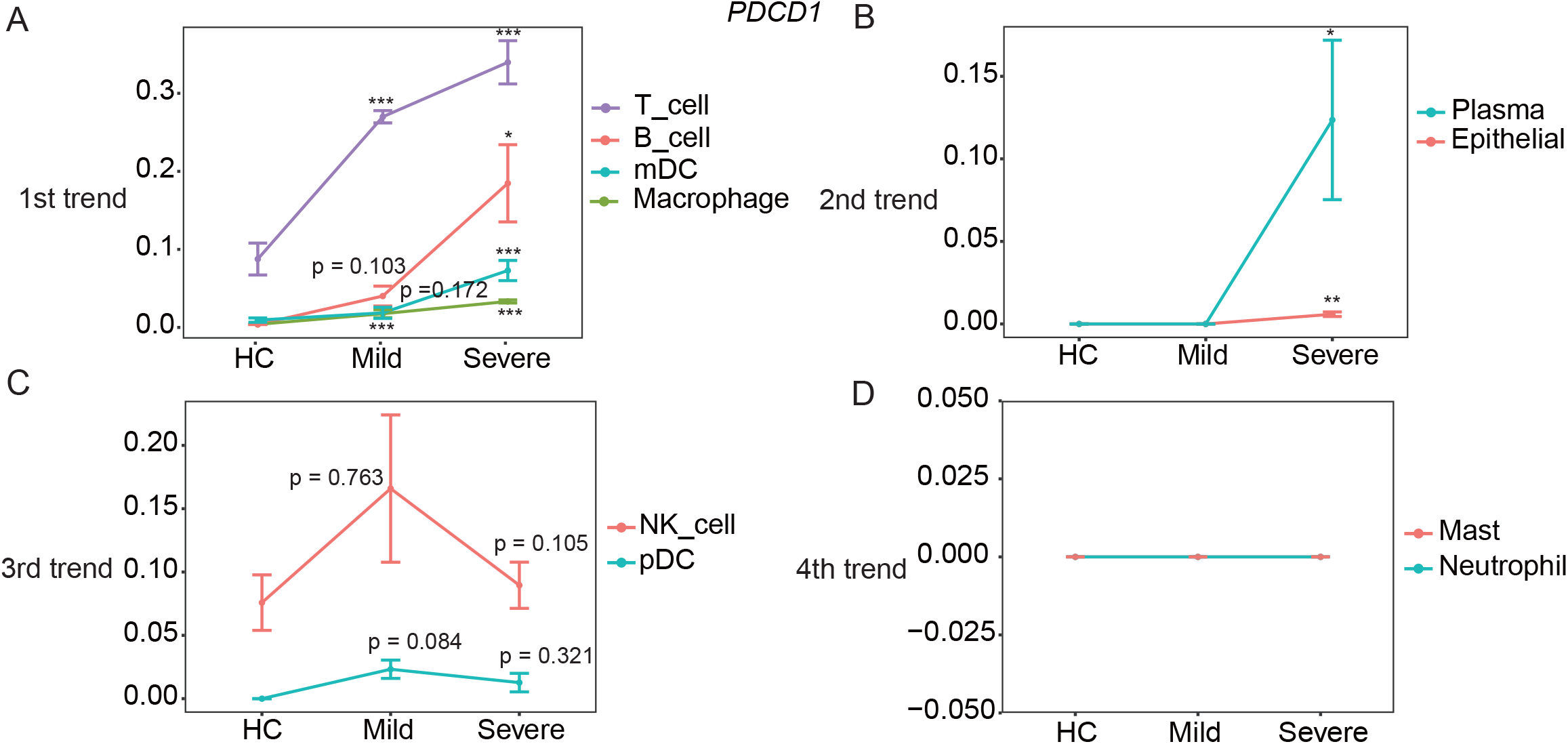

The *CD274* expression in macrophages, mast cells, pDC, and T cells (1^st^ trend) correlated well with COVID-19 severity (Fig. 3A) and was specifically increased in plasma cells of severe patients (2^nd^ trend) (Fig. 3B). However, in neutrophil, mDC, B cells and NK cells, the expression of *CD274* was upregulated in patients with mild COVID-19, then decreased in severe patients (Fig. 3C). In contrary to the 3^rd^ trend, the expression of *CD274* was decreased in mild patients but increased in severe patients (Fig. 3D).

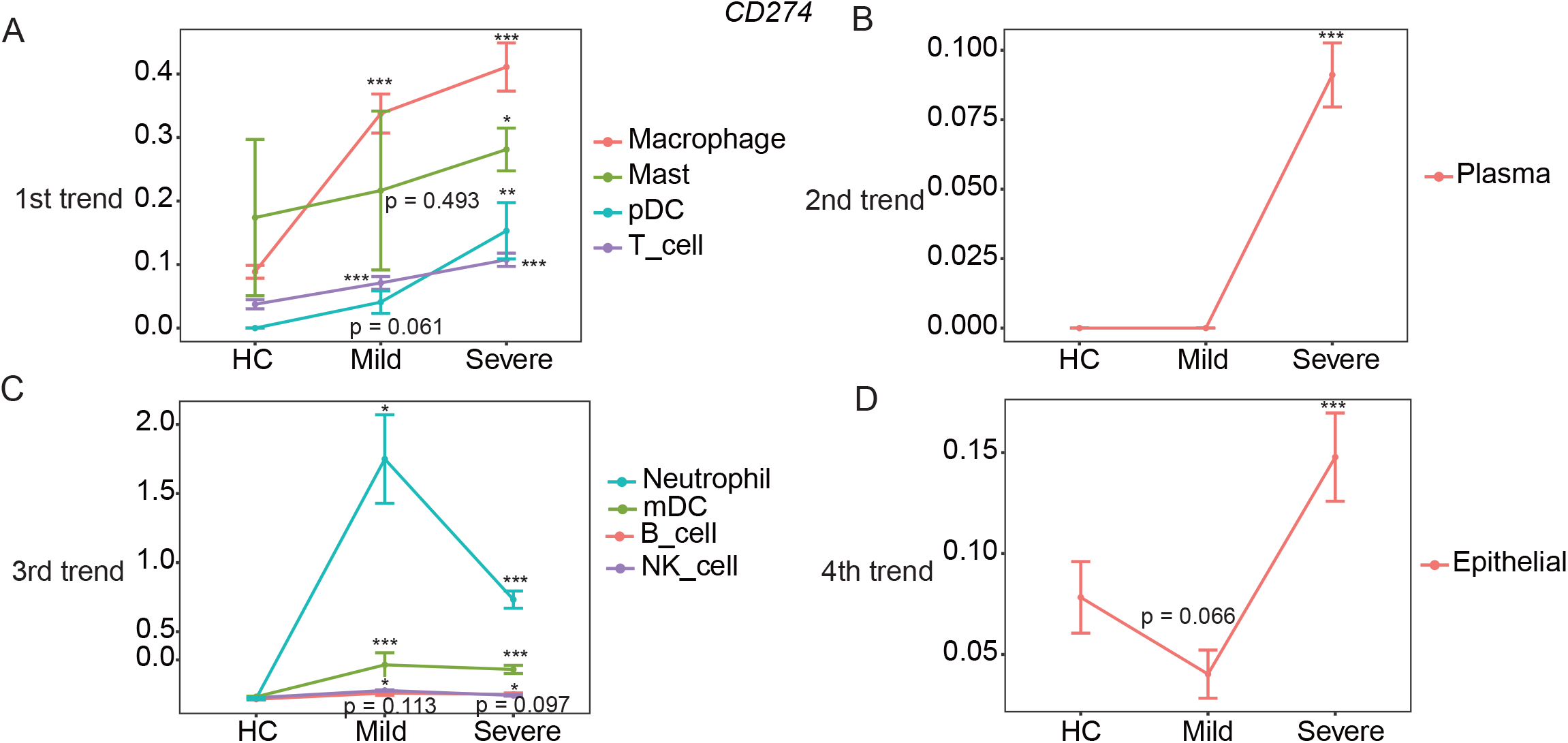

When analyzed at the individual level, expression of *PDCD1* and *CD274* was also elevated in mild and severe patients (Fig. 4A, 4B). Overall, *PDCD1* expression in T cells, B cells, mDCs, and macrophages and *CD274* expression in macrophages, mast cells, pDC, and T cells correlated well with COVID-19 severity. Furthermore, *PDCD1* and *CD274* expression was specifically increased in epithelial and plasma cells of severe patients.

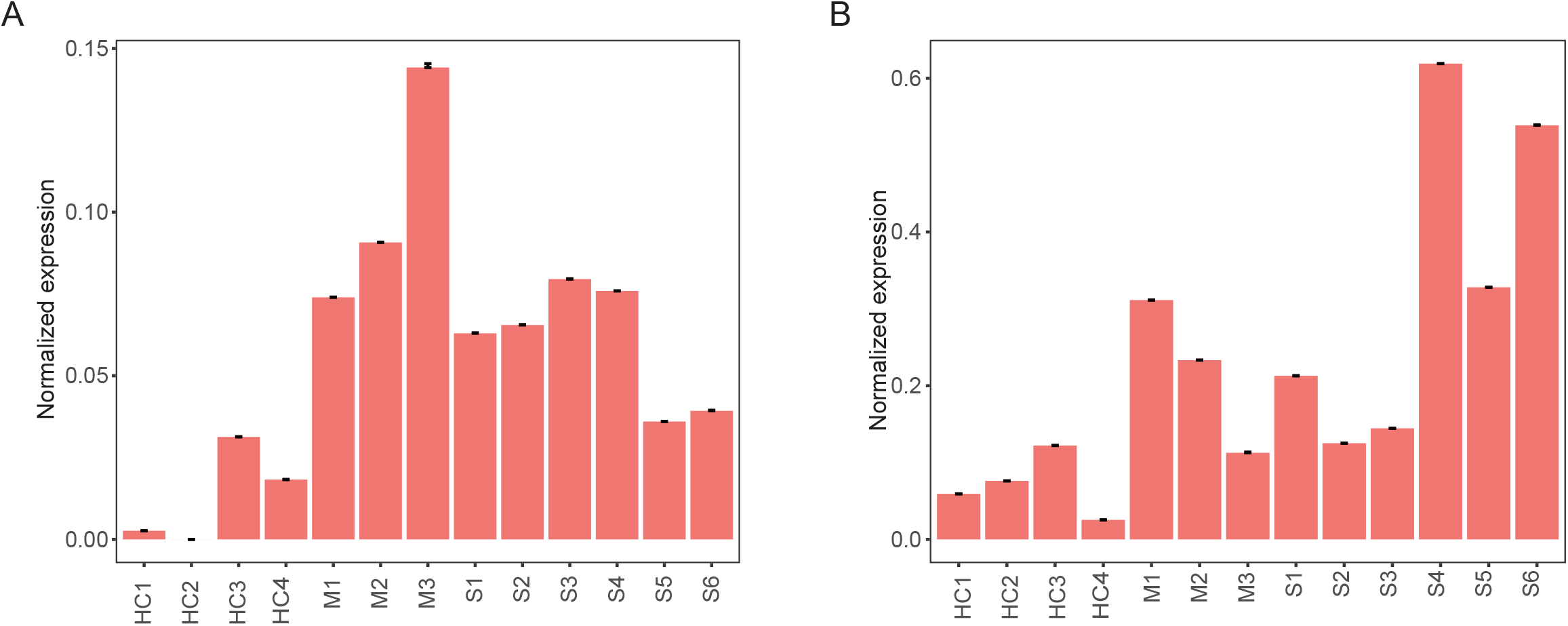

### The expression of *STAT1* was increased in patients with mild and severe COVID-19

Inflammatory signaling participates in modulating PD-L1 expression, particularly, STAT1, which can be activated by IFNγ or interleukin 6 (IL-6), is a crucial regulator for PD-L1 expression [7, 8]. Furthermore, plasma IFNγ level [3] and the IL-6 level in bronchoalveolar lavage fluid (BALF) [6] were reported to be increased in COVID-19 patients. Consistently, *STAT1* was found upregulated in both mild and severe patients (Fig. 5), suggesting increased *CD274* expression might at least partly resulting from increased *STAT1* level in COVID-19 patients.

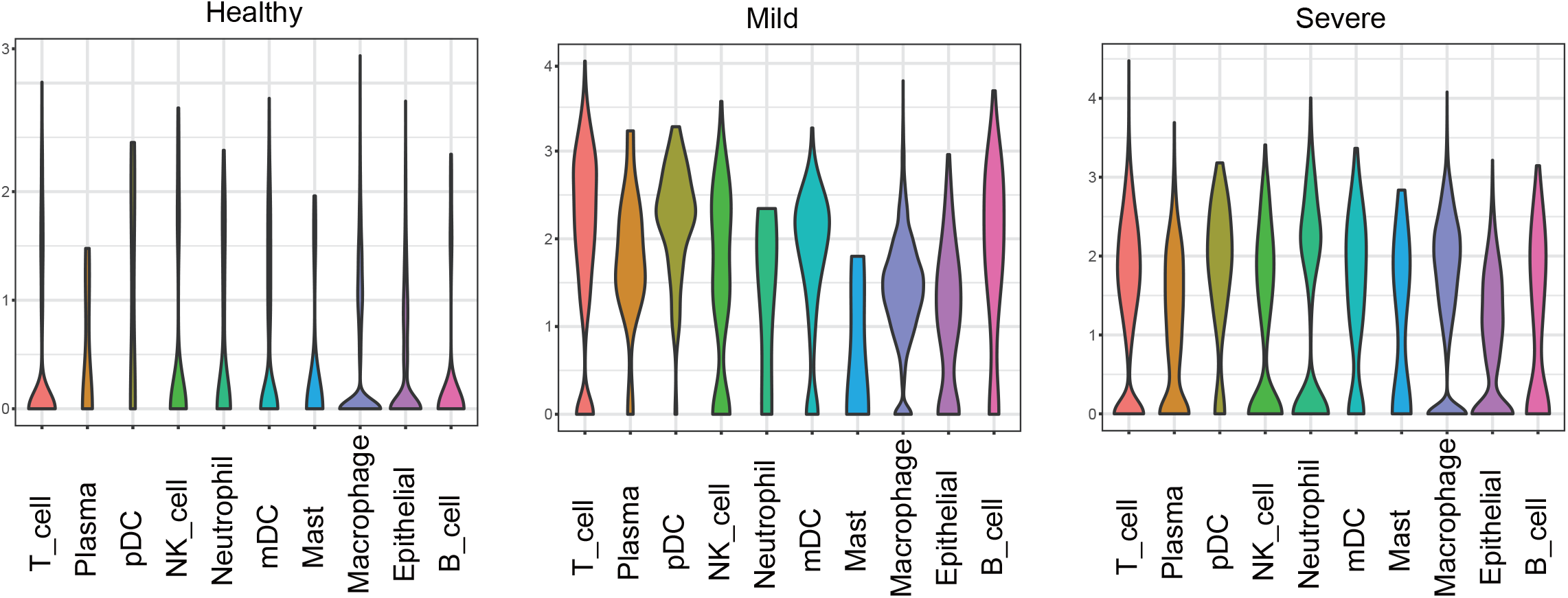

### The landscape of immune checkpoint molecules in COVID-19 patients

To further elucidate the immune checkpoint landscape of COVID-19 patients, expression of classical inhibitory and stimulatory checkpoint molecules was assessed. For inhibitory checkpoint molecules (ICMs), expression of *CD160*, *CD244*, *PDCD1*, *BTLA*, *TIGIT*, *LAG3*, *KLRG1*, and *ADORA2A* were increased in mild patients compared to HC and severe patients while expression of *CTLA4*, *HAVCR2*, *IDO1*, and *CD276* were highest in severe patients (Fig. 6A). Regarding stimulatory checkpoint molecules (SCMs), expression of *TNFRSF9*, *CD28*, *ICOS*, and *CD27* was elevated in mild patients in comparison to HC and severe patients while expression of *TNFRSF18* and *CD40* were highest in severe patients (Fig. 6B).

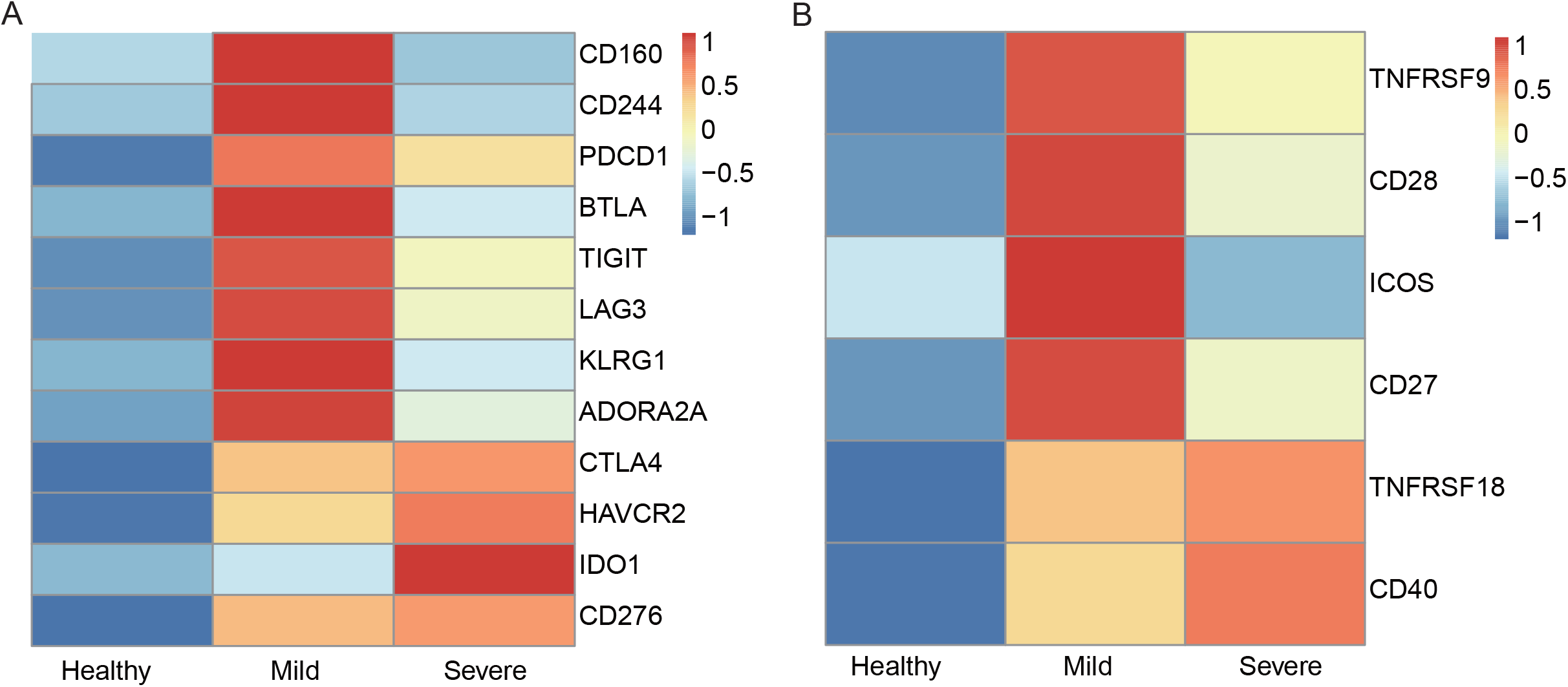

When analyzed at the cell subpopulation level, unique expression patterns of ICMs and SCMs were demonstrated in each cell subpopulation (Fig. 7, Fig. 8). Expression patterns of ICMs were different between mild and severe patients (Fig. 7). Most categories and the highest intensity of ICMs were enriched in severe patients, except in T cells and NK cells. The results indicated that the immune cells might be more exhausted in severe patients with COVID-19 compared to healthy people and even mild patients. In terms of SCMs, macrophages, mast cells, and pDC of severe patients got more enriched than that of mild patients and healthy donors (Fig. 8). In T cell population, more ICMs and SCMs were enriched in patients with mild and severe COVID-19 than that of healthy donors, but there was no dramatic difference between mild and severe patients themselves (Fig. 7, Fig. 8).

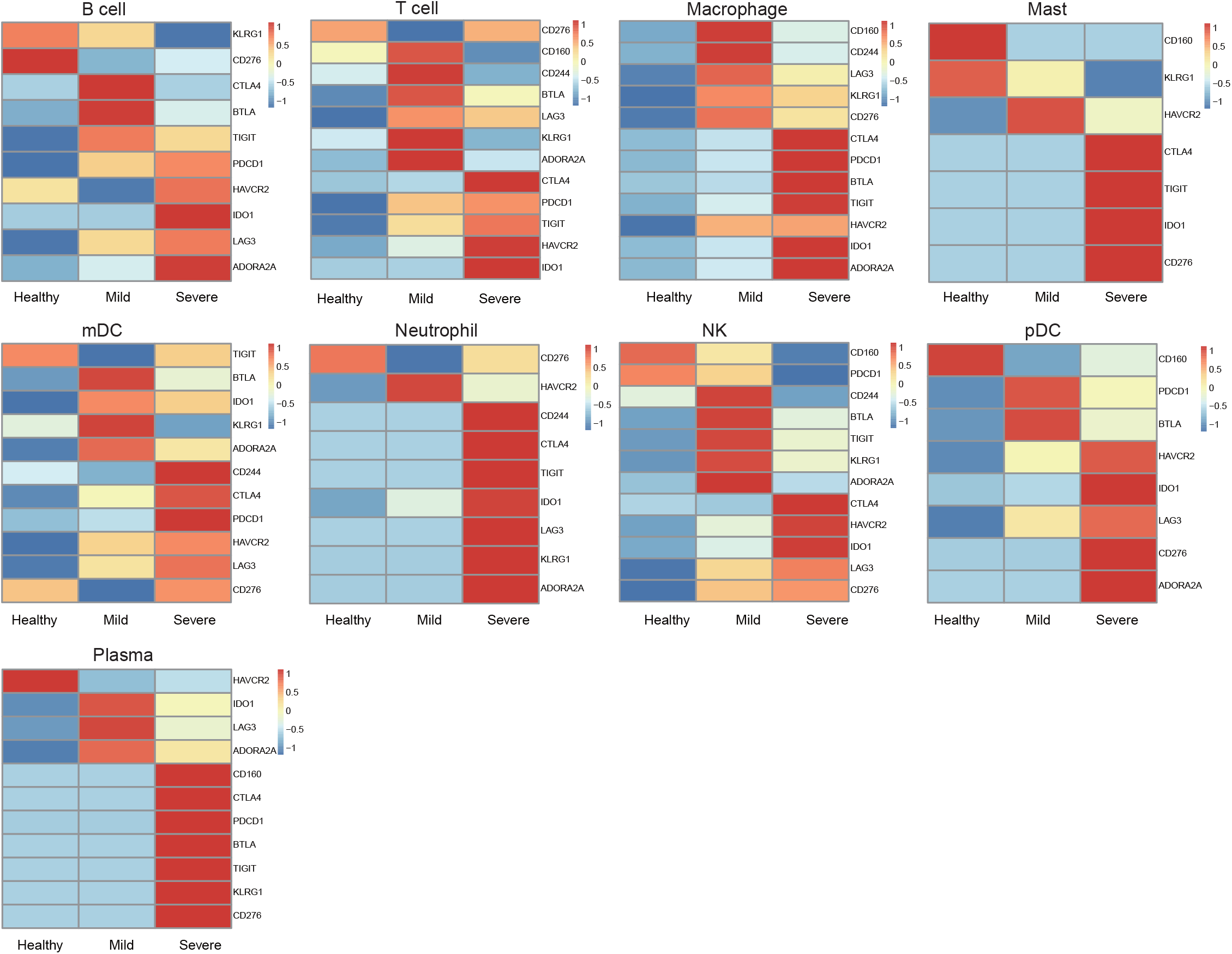

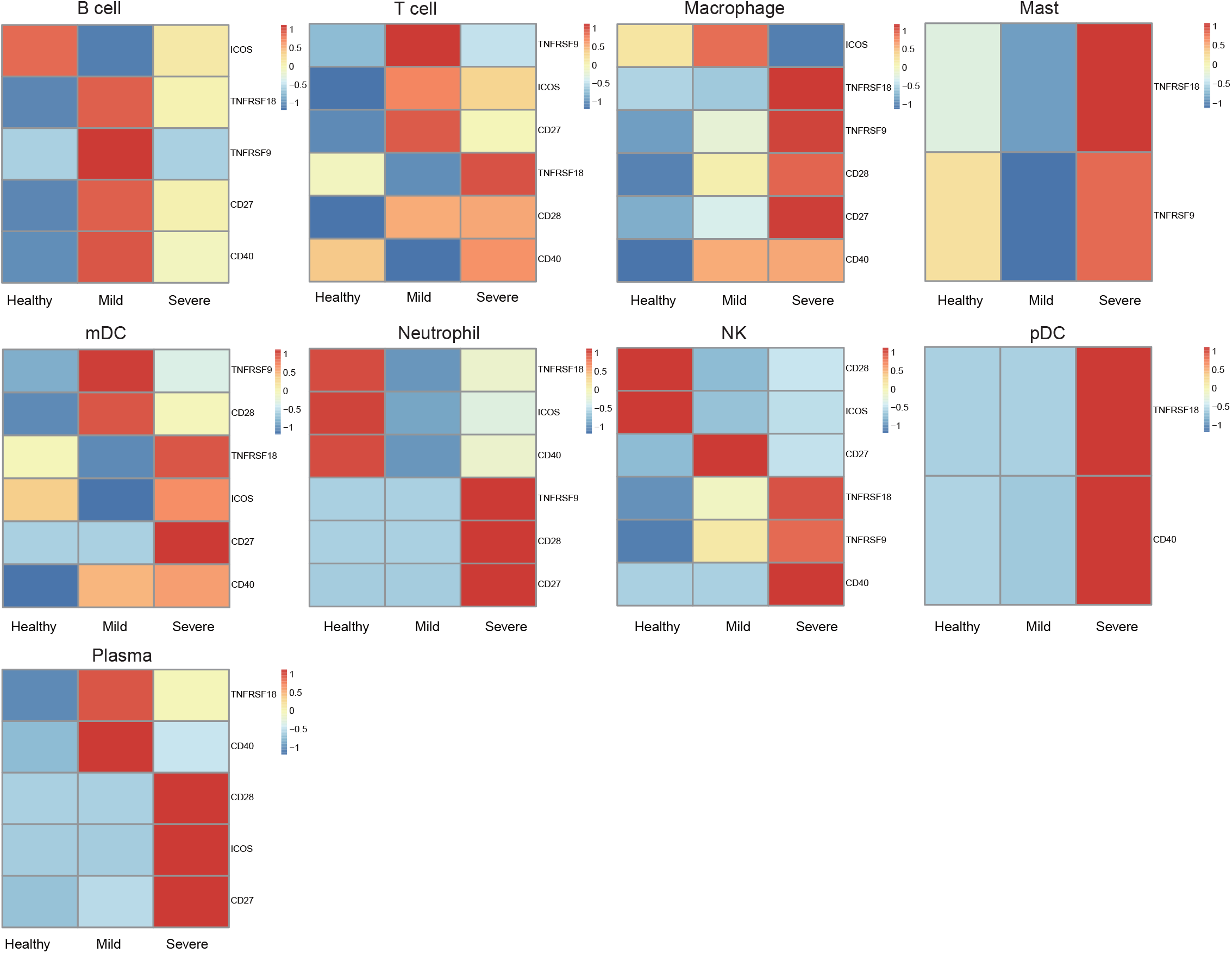

## Discussion

Since COVID-19 is pandemic and threatening thousands of people’s life, it is urgent and essential to investigate the molecular mechanism of the immune pathogenesis of the disease. Compared to healthy controls, *PDCD1* expression in T cells, B cells, mDCs, and macrophages (Fig. 2) and *CD274* expression in macrophages, mast cells, pDCs, and T cells (Fig. 3) were upregulated in COVID-19 patients, and correlated well with COVID-19 severity. Moreover, expression of *PDCD1* and *CD274* was specifically increased in plasma cells of severe patients (Fig. 2, 3), which could serve as a biomarker for prognosing the severity of COVID-19. Many clinical trials for treating COVID-19 are ongoing. Among them, one clinical trial uses PD-1 monoclonal antibody to block PD-1 in COVID-19 patients (NCT04268537). Based on our results, *PDCD1* expression was dramatically upregulated in T cells and macrophages especially in severe patients (Fig. 2) and its blockade would further increase the secretion of multiple pro-inflammatory cytokines, which will enhance the cytokine release syndrome reported in COVID-19 patients and possibly associated with disease severity [2, 3], leading to further tissue damage or even more death especially in severe COVID-19 patients [9, 10]. A current study supports that checkpoint inhibitor immunotherapy is risky for severe outcomes in SARS-CoV-2-infected cancer patients, though these patients were treated with immune checkpoint inhibitors (ICI) before SARS-CoV-2 infection [11].

## Conclusion

Our research provides valuable information about COVID-19 and its severity correlated well with the expression of *PDCD1* and *CD274*.

## Author contributions

Gao Q. supervised the project, designed the bioinformatic analysis, wrote and revised the manuscript. Liu S. performed the bioinformatic analysis. Ding R. dealt with the data. Chen H., Dong X., Xie J., Li Y., Chen L., and Liu H. helped with the data management.

## Acknowledgments

We sincerely thank the support provided by China National GeneBank and this research was supported by the Guangdong Enterprise Key Laboratory of Human Disease Genomics (2020B1212070028), and Science, Technology and Innovation Commission of Shenzhen Municipality (JSGG20180508152912700).

## References

1. Wilk A.J., Rustagi A., Zhao N.Q., Roque J., Martínez-Colón G.J., McKechnie J.L., et al., A single-cell atlas of the peripheral immune response in patients with severe COVID-19. Nat Med, 2020. 26: p. 1070–76.

2. Chen Nanshan, Zhou Min, Dong Xuan, Qu Jieming, Gong Fengyun, Han Yang, et al., Epidemiological and clinical characteristics of 99 cases of 2019 novel coronavirus pneumonia in Wuhan, China: a descriptive study. The Lancet, 2020. 395(10223): p. 507–513.

3. Huang Chaolin, Wang Yeming, Li Xingwang, Ren Lili, Zhao Jianping, Hu Yi, et al., Clinical features of patients infected with 2019 novel coronavirus in Wuhan, China. The Lancet, 2020. 395(10223): p. 497–506.

4. Liu Y., Zhang C., Huang F., Yang Y., Wang F., Yuan J., et al., 2019-novel coronavirus (2019-nCoV) infections trigger an exaggerated cytokine response aggravating lung injury. ChinaXiv:202002.00018, 2020.

5. Qin Weiting, Hu Lipeng, Zhang Xueli, Jiang Shuheng, Li Jun, Zhang Zhigang, et al., The Diverse Function of PD-1/PD-L Pathway Beyond Cancer. Frontiers in Immunology, 2019. 10.

6. Liao M., Liu Y., Yuan J., Wen Y., Xu G., Zhao J., et al., Single-cell landscape of bronchoalveolar immune cells in patients with COVID-19. Nat Med, 2020.

7. Garcia-Diaz A., Shin D. S., Moreno B. H., Saco J., Escuin-Ordinas H., Rodriguez G. A., et al., Interferon Receptor Signaling Pathways Regulating PD-L1 and PD-L2 Expression. Cell Rep, 2017. 19(6): p. 1189–1201.

8. Moon J. W., Kong S. K., Kim B. S., Kim H. J., Lim H., Noh K., et al., IFNgamma induces PD-L1 overexpression by JAK2/STAT1/IRF-1 signaling in EBV-positive gastric carcinoma. Sci Rep, 2017. 7(1): p. 17810.

9. Zhang C., Wu Z., Li J. W., Zhao H.,Wang G. Q., Cytokine release syndrome in severe COVID-19: interleukin-6 receptor antagonist tocilizumab may be the key to reduce mortality. Int J Antimicrob Agents, 2020. 55(5): p. 105954.

10. Chua Robert Lorenz, Lukassen Soeren, Trump Saskia, Hennig Bianca P., Wendisch Daniel, Pott Fabian, et al., COVID-19 severity correlates with airway epithelium–immune cell interactions identified by single-cell analysis. Nat Biotechnol, 2020.

11. Robilotti E. V., Babady N. E., Mead P. A., Rolling T., Perez-Johnston R., Bernardes M., et al., Determinants of COVID-19 disease severity in patients with cancer. Nat Med, 2020.

